# SecretoMyc, a web-based database on mycobacteria secreted proteins and structure-based homology identification using bio-informatics tools

**DOI:** 10.1101/2023.04.21.537767

**Authors:** Jérôme Gracy, Katherine Vallejos-Sanchez, Martin Cohen-Gonsaud

## Abstract

To better understand the interaction between the host and the *Mycobacterium tuberculosis* pathogen, it is critical to identify its potential secreted proteins. While various experimental methods have been successful in identifying proteins under specific culture conditions, they have not provided a comprehensive characterisation of the secreted proteome. We utilized a combination of bioinformatics servers and in-house software to identify all potentially secreted proteins from six mycobacterial genomes through the three secretion systems: SEC, TAT, and T7SS. The results are presented in a database that can be crossed with selected proteomics and transcriptomics studies (https://secretomyc.cbs.cnrs.fr/myc). In addition, thanks to the recent availability of Alphafold models, we developed a tool in order to identify the structural homologues among the mycobacterial genomes.

---

Mycobacterial infection relied on a multiple of immune cell response perturbation, from the phagosome maturation to the cytokines secretion^1^. The response modification is dependent of a multitude of effectors either lipidic or protein molecules. It is estimated that over 20% of bacterial proteins have functions outside the bacterial cytoplasm and are exported to their designated locations by protein export systems^2^. The functions of the exported proteins are essential for physiological processes (*i*.*e*. the cell-wall maintenance) but also, in the case of pathogenic bacteria they are crucial for virulence. To comprehend the interplay between host and pathogens it is then essential to identify the putative secreted proteins. Biochemical, genetic and imaging tools have been developed to evaluate protein secretion^3^. If reported-based assay^4^ and functional screens^5^ have been used, but mass spectrometry is the most commonly employed technique in order to identify secreted or cell-wall associated protein in various strains or culture conditions.

Proteomics studies have demonstrated their ability to identify proteins in specific growth conditions^6–8^. While these studies can reveal the secretion patterns in a particular condition, they are not designed to identify the entire bacterial secretome. It is unsurprising to observe variations in secretion patterns across different bacterial experiments. As a consequence, data from various proteomic studies on secreted mycobacterial proteins have shown a weak overlap for proteins identified as secreted. Furthermore, the host cell environment also plays an important role, as recently revealed by two studies focusing on the identification of secreted proteins during infection^4,9^.

*M. tuberculosis* possesses three different secretion systems^10^. The general secretion (Sec) and the twin-arginine translocation (Tat) pathways perform the bulk of protein export and are both essential. Proteins exported by the Sec pathway are distinguished by the presence of an N-terminal signal recognized by the SecA protein before translocation. The Tat pathway exports preproteins containing N-terminal signal peptides with a twin-arginine motif for binding to the TatC protein. *M. tuberculosis* has also specialized export pathways that transport specific subsets of proteins. Five specialized ESX export systems (ESX-1 to ESX-5) are present in *M. tuberculosis* with some of them essential for virulence^11^. Although the ESX systems were first discovered in *M. tuberculosis*, they also exist in a few other Gram-positive bacteria. The ESX systems are also referred to as Type VII secretion systems (T7SS). Proteins secreted by T7SS lack Sec or Tat signal peptides, instead secretion relies on a combination of sequence and structural motifs. Based on the identification of various T7SS secreted proteins, two secretion motifs (YxxxD/E and WxG) included in a flexible loop that participates in a helix-turn-helix structure^12,13^ were identified. Two proteins with one each of the motifs interacts are exported as an heterodimer after binding to the intracellular chaperone EspG protein^14^.

Here we used and combines multiple bioinformatics server and in-house tools in order to identify all the putative secreted mycobacterial proteins. We analyzed the 3906 *M. tuberculosis* H37Rv sequences using an in-house pipelining tool for large-scale integrative bioinformatics^15^ (PAT pipeline). First, known signal peptides and/or structural features necessary for secretion were predicted using dedicated available software^16,17^ (*i*.*e*. SignalP v4.1 and PredTAT). In addition, transmembrane segments were inferred using either Uniprot annotations or the TmHMM prediction software^18^, then we also looked at the number of predicted transmembrane segments and the position of the last transmembrane segment to identify signals potentially missed by the various servers and to avoid confusion with membrane proteins. For the T7SS, based on previous work we aim to identified the combination of sequence and structural motifs. We selected the *M. tuberculosis* proteins whose Alphafold models obtained from the EBI database (https://alphafold.ebi.ac.uk/) had at least 70 residues in helical conformation in the 100 first positions, and whose sequences had a WxG motif between positions 30 and 79 or a YxxxD/E motif between positions 80 and 99. This Alphafold-based selection method detected 108 putative T7SS proteins in *M. tuberculosis* (alignments available at (https://secretomyc.cbs.cnrs.fr/myc/myctu_H70-99_PE0-19_Wxg30-79_or_YDE80-99.htm).

These data were crossed with various data from proteomics and transcriptomics studies and five additional mycobacterial genomes (*M. marinum, M. bovis, M. abscessus, M*. avium and *M. smegmatis*) underwent the same process to compare predictions between various mycobacterial species, specifically in regards to their ability to survive within infected cells. Thus, we identified 176 T7SS, 79 Tat, and 462 Sec *M. tuberculosis* proteins with characteristic secretion signals. The data are presented in web-page at this address: https://secretomyc.cbs.cnrs.fr/myc

Next, we first applied the homology modeling procedure to identify distant homologues of the proteins, and the resulting information was used to exclude false positive proteins. For instance, some proteins belonging to the TerR cytoplasmic transcriptional regulator family possess sequence and structural features matching the T7SS secretion motif but are not secreted, and as consequence excluded from our secretome. We took advantages of the breakthrough brought by AlphaFold^19^ to insert in our database the generated models. Those models can be used to detect structural homologues and then correct and/or complement the current genomic annotations as well as it provides information on domain delimitation. Here, we used the models to identify homologues within the genome. We first collected a structure database including all Alphafold models of the *M. tuberculosis* proteome in the EBI database (https://alphafold.ebi.ac.uk/) and a representative set of protein domain structures obtained from the SCOP classification database^20^. Structure pairs were structurally aligned using TMalign^21^ and the pairs sharing a TM-score above 0.55 were then hierarchically aggregated yielding structural clusters which we called AlphaClans. When a *M. tuberculosis* Alphafold model belongs to an AlphaClan, its member structures are listed at the bottom of the corresponding *M. tuberculosis* report, with pairwise TM-scores, RMSD, and sequence identity percentages. The secretion scores which are also listed for each AlphaClan member can help to better assess if the considered *M. tuberculosis* protein is secreted or not.

The home page of the web server provides access to the SecretoMyc database through a global table where information on all *M. tuberculosis* proteins is synthetized: database cross-references, homology searches, secretion predictions, structural models, domain families and proteomics experiments (Figure 1). The display of table columns can be customized using the left panel with toggle buttons. Column data can be sorted or searched using simple or advanced query form.

**Figure 1:**
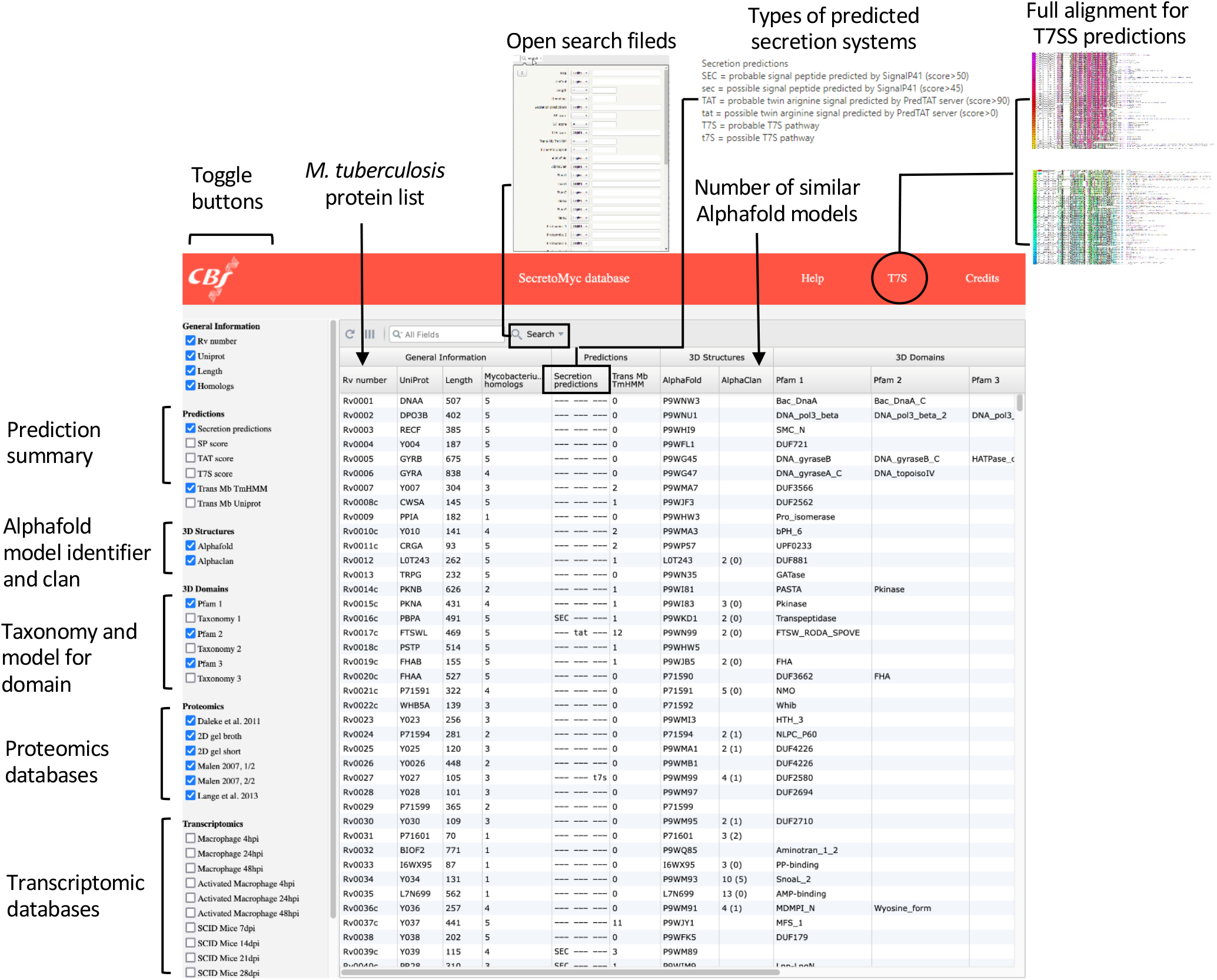
The home page of the web server that provides access to the SecretoMyc prediction database is an interactive analysis toolkit. The home page displays the all *M. tuberculosis* proteome with quick access to essentials information: Secretion prediction, taxonomy and domain identification, structural homologue using the AlphaClan tool and homologues within other mycobacterial genomes (*M*.*Marinum, M*.*bovis, M*.*abscessus, M*.*avium*, and *M*.*smegmatis*) as well as results from proteomics and transcriptomics studies. The columns are sortable, and the user can utilize an open search field to select a portion of the secretome based on one or multiple criteria from the available cross-referenced information.

Clicking on a table line will pop up a detailed report of all cross-references, homologues, predictions, and classifications obtained on the corresponding protein (Figure 2). This report provides many important information for analyzing the protein: alignment with orthologs from close genomes, synthetic table with transmembrane and secretion predictors, list of similar Pfam families, PDB structures or Alphafold models.

**Figure 2:**
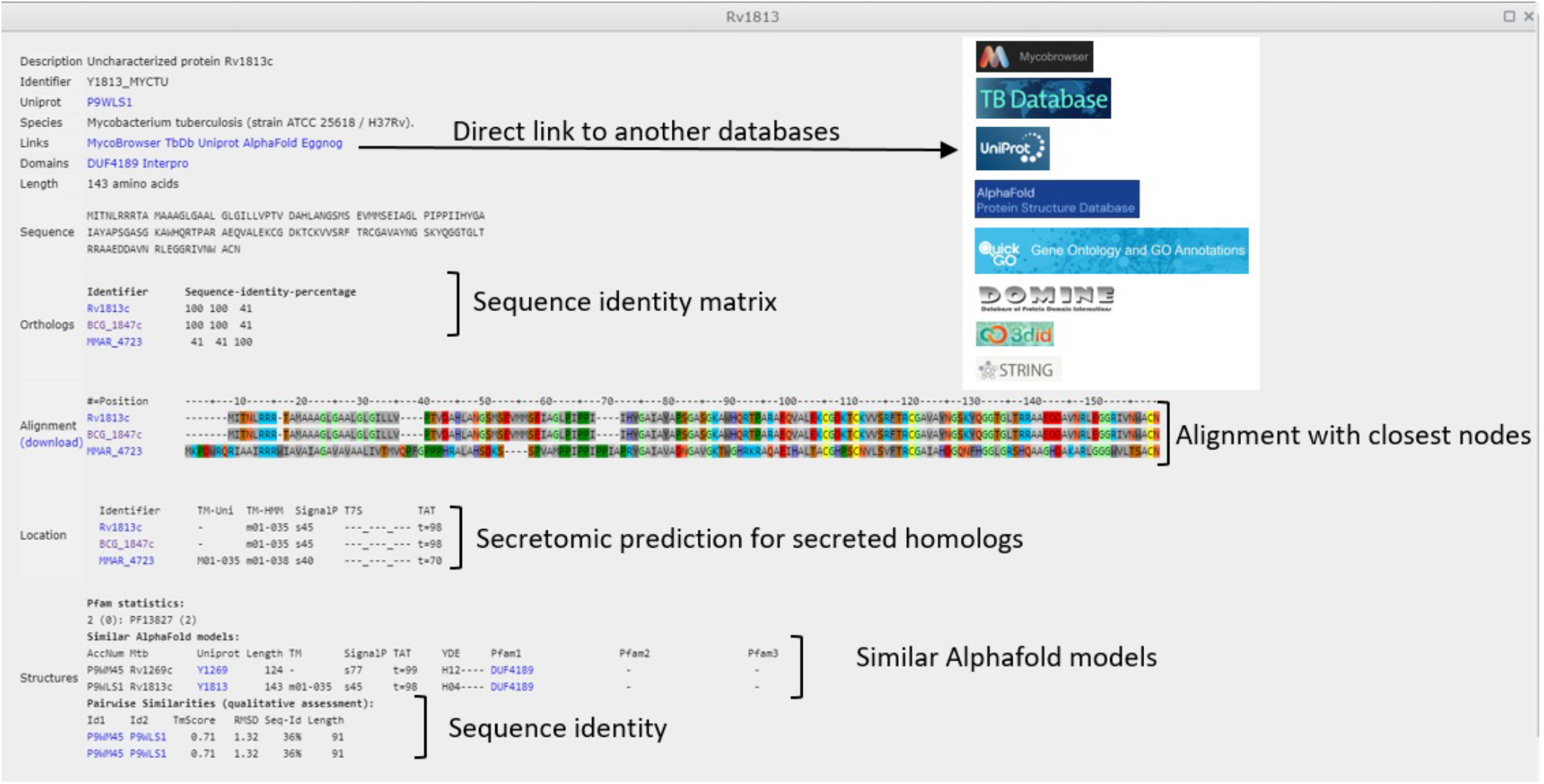
Clicking on a corresponding line from the homepage will lead to a protein page that contains all available information about the protein, including a description and direct links to major protein databases, protein sequence, protein homolog identity matrix and alignments, secretion predictions for all homologues, and structural homology prediction using Alphaclan.

In summary, the database provides a convenient way for users to determine whether a protein of interest may be secreted, and to identify homologues among six mycobacterial proteomes, as well as more distant structural homologues using the available Alphafold models and AlphaClan classification. Each individual page presents all the required information for evaluating the results. Furthermore, the database facilitates the integration of data from proteomics and transcriptomics studies, allowing users to set thresholds for selecting putatively secreted proteins based on their corresponding RNA expression levels. While the database currently includes only a few selected studies, more can be incorporated upon request from users.

## Material and methods

Six mycobacteria proteomes (*M. tuberculosis, M. bovis, M. avium, M. marinum, M. smegmatis* and *M. abcessus*) were retrieved using the “Proteome” server of the Uniprot database^22^. Secretion pathways, structural predictions, domain architectures, proteomics and transcriptomics data were predicted or collected for each of these proteins. The resulting information was summarized in individual entries describing each *M. tuberculosis* protein. The whole SecretoMyc database is accessible through an interactive web table whose columns can be customized, sorted and queried.

More precisely, each protein information was obtained in the following way: General secretion pathway (SEC) peptide signals were predicted using the program SignalP version 4.1^16^ with model trained on Gram-positive bacteria and D-cutoff 0.45. Twing-arginine secretion pathway (TAT) signals were predicted using the web server PRED-TAT (http://www.compgen.org/tools/PRED-TAT)^17^.

Type VII secretion pathway (T7SS) proteins: A sequence database composed of *M. tuberculosis* protein fragments was built in the following way: the sequence fragments were composed of the 110 first residues of each *M*.*tuberculosis* protein, the identifier of each protein fragment was obtained by concatenating the Tuberculist identifier^23^, the gene name, the first PFAM domain^24^, the number of residues in helix conformations, and the starting positions of the identified motifs WxG and YxxxD/E when they were found. This protein fragment database was searched using seven iterations of Hmmer (http://hmmer.org/) with E-value cutoff 1 and a 50% query overlap cutoff using as input query the alignment of all ESX proteins which are all known to belong to T7SS. The full sequences corresponding to the hits detected by Hmmer were then aligned using MAFFT, resulting in 187 aligned T7S proteins with gene names, PFAM domains, helix counts, and PPE, WxG, YxxxD/E annotations. Finally, for each aligned *M. tuberculosis* hit were collected all interacting proteins with a combined score above 800 according to the STRING database^25^.

Transmembrane proteins were predicted using the TmHMM software. The transmembrane features were also retrieved from the Uniprot database^22^. The number of predicted or featured transmembrane segments were stored in the SecretoMyc database for each *M. tuberculosis* protein.

Alphafold 3D models computed for the whole *M. tuberculosis* proteome were obtained from the EBI web server (https://alphafold.ebi.ac.uk). A database of 3D structures was then built by merging these 3D *M. tuberculosis* models with a list of representative experimental domain structures obtained from the SCOP database filtered at 40% maximum sequence identity. All these 3D structures were hierarchically aggregated according to a similarity score combining the BLOSUM62 substitution score^26^ of their aligned sequences and the similarity of their secondary structure composition. This initial classification step resulted in a binary tree whose nodes were checked following a bottom-up tree scan. For each node, a protein pair representative of its 2 descending branches was selected and compared using the structural alignment program TM-align^21^. Each branch representative was the protein closest to the consensus sequence of the protein cluster under the considered node branch. If the structural TM score was above 0.55, the checked node was validated and its father node was further inspected. Otherwise, the leaf cluster under each branch of the non-validated node was saved as an Alphafold clan of similar protein structures sharing Tm-scores above 0.55. The tree traversal was then skipped to the next non checked tree-bottom leaf.

3D domains were obtained from the PFAM database and up to 3 domains were stored in our database for each *M. tuberculosis* protein.

Furthermore, we added data proteomics research to enable the user to compare our findings with proteomics studies. *M. tuberculosis* culture filtrated protein identify after mass spectrometry analysis focused pI range 4.0-4.7 and the Mr range 6-20kDa of the 2-DE pattern were added^27^ (33 proteins identified), as well two previous studies and more general characterisation after short and long culture from the same group that were previously available on MPIIB web-page (138 and 33 proteins identified). Similar studies with different results were also added^6^ (159 and 254 proteins identified). Data from a specific bioinformatics study focused on T7SS was also added^13^ (92 proteins identified).

In the context of the host pathogen interaction and immune response perturbation, it is also interesting for the user to be able to compare our data with transcriptomics data obtained during infection. We added two sets of data of genes expressed differentially as a consequence of intraphagosomal residence^28^ (compared to broth culture) with or without activation at three time points (4, 24 and 48 hours post infection) (Schnappinger2003) and transcription profile of genes expressed during the course of early tuberculosis in immune-competent (BALB/c) and severe combined immune-deficient (SCID) hosts in comparison with growth in medium at three time points (7, 14 and 21 days post infection).

All this computed information was compiled in a global database called SecretoMyc. It is accessible through a web server (http://secretomyc.cbs.cnrs.fr) with a Javascript interface based on W2UI (https://w2ui.com/web/). For easier use, the displayed columns of the global table shown in the server home page can be selected using the dedicated left panel. The protein entries can also be sorted according to each column and a searching form permits the construction of composite queries for selecting protein subsets.

For each *M. tuberculosis* protein, an individual entry was stored in the SecretoMyc database by compiling the following information:

- Uniprot identifier and accession number, gene name, species name, sequence length, amino acid sequence.
- Links to complementary databases Uniprot (https://www.uniprot.org/), TBDB (http://tbdb.bu.edu/), Mycobrowser (https://mycobrowser.epfl.ch/), AlphaFold (https://alphafold.ebi.ac.uk/), GO (http://geneontology.org/), EggNog (http://eggnog5.embl.de/), 3Did (https://3did.irbbarcelona.org/), Domine (https://manticore.niehs.nih.gov/cgi-bin/Domine) and String (https://string-db.org/).
- Orthologous proteins detected in the 5 close mycobacteira genomes (*M. bovis, M. avium, M. marinum, M. smegmatis* and *M. abcessus*). The ortholog groups were clustered using the Kclust program^29^.
- Alignment of all orthologs built using the multiple sequence alignment program MAFFT^30^.
- Secretion pathway and transmembrane annotations are summarized in global “Location” table listing all ortholog predictions.
- Similar SCOP domains and Alphafold models detected for each protein as explained above. The detected pairwise 3D similarities are assessed using the various scores provided by the structural alignment program TM-align (TM-score, RMS deviation, Sequence identity percentage, Alignment length).

## Notes

https://secretomyc.cbs.cnrs.fr/myc

### Competing Interest Statement

The authors have declared no competing interest.

## Bibliography

1. Pai, M., Behr, M.A., Dowdy, D., Dheda, K., Divangahi, M., Boehme, C.C., Ginsberg, A., Swaminathan, S., Spigelman, M., Getahun, H., et al. (2016). Tuberculosis. Nat Rev Dis Primers 2, 1–23. 10.1038/nrdp.2016.76.

2. Kostakioti, M., Newman, C.L., Thanassi, D.G., and Stathopoulos, C. (2005). Mechanisms of Protein Export across the Bacterial Outer Membrane. J Bacteriol 187, 4306–4314. 10.1128/JB.187.13.4306-4314.2005.

3. Maffei, B., Francetic, O., and Subtil, A. (2017). Tracking Proteins Secreted by Bacteria: What’s in the Toolbox? Front Cell Infect Microbiol 7, 221. 10.3389/fcimb.2017.00221.

4. Perkowski, E.F., Zulauf, K.E., Weerakoon, D., Hayden, J.D., Ioerger, T.R., Oreper, D., Gomez, S.M., Sacchettini, J.C., and Braunstein, M. (2017). The EXIT Strategy: an Approach for Identifying Bacterial Proteins Exported during Host Infection. mBio 8.p 10.1128/mBio.00333-17.

5. Heidtman, M., Chen, E.J., Moy, M.-Y., and Isberg, R.R. (2009). Large scale identification of Legionella pneumophila Dot/Icm substrates that modulate host cell vesicle trafficking pathways. Cell Microbiol 11, 230–248. 10.1111/j.1462-5822.2008.01249.x.

6. Målen, H., Berven, F.S., Fladmark, K.E., and Wiker, H.G. (2007). Comprehensive analysis of exported proteins from Mycobacterium tuberculosis H37Rv. Proteomics 7, 1702–1718. 10.1002/pmic.200600853.

7. de Souza, G.A., Leversen, N.A., Målen, H., and Wiker, H.G. (2011). Bacterial proteins with cleaved or uncleaved signal peptides of the general secretory pathway. J Proteomics 75, 502–510. 10.1016/j.jprot.2011.08.016.

8. Albrethsen, J., Agner, J., Piersma, S.R., Højrup, P., Pham, T.V., Weldingh, K., Jimenez, C.R., Andersen, P., and Rosenkrands, I. (2013). Proteomic profiling of Mycobacterium tuberculosis identifies nutrient-starvation-responsive toxin-antitoxin systems. Mol Cell Proteomics 12, 1180–1191. 10.1074/mcp.M112.018846.

9. Penn, B.H., Netter, Z., Johnson, J.R., Von Dollen, J., Jang, G.M., Johnson, T., Ohol, Y.M., Maher, C., Bell, S.L., Geiger, K., et al. (2018). An Mtb-Human Protein-Protein Interaction Map Identifies a Switch between Host Antiviral and Antibacterial Responses. Mol Cell 71, 637–648.e5. 10.1016/j.molcel.2018.07.010.

10. Feltcher, M.E., Sullivan, J.T., and Braunstein, M. (2010). Protein export systems of Mycobacterium tuberculosis: novel targets for drug development? Future Microbiol 5, 1581–1597. 10.2217/fmb.10.112.

11. Vaziri, F., and Brosch, R. (2019). ESX/Type VII Secretion Systems-An Important Way Out for Mycobacterial Proteins. Microbiol Spectr 7.p 10.1128/microbiolspec.PSIB-0029-2019.

12. Pallen, M.J., Chaudhuri, R.R., and Henderson, I.R. (2003). Genomic analysis of secretion systems. Curr Opin Microbiol 6, 519–527. 10.1016/j.mib.2003.09.005.

13. Daleke, M.H., Ummels, R., Bawono, P., Heringa, J., Vandenbroucke-Grauls, C.M.J.E., Luirink, J., and Bitter, W. (2012). General secretion signal for the mycobacterial type VII secretion pathway. Proc Natl Acad Sci U S A 109, 11342– 11347. 10.1073/pnas.1119453109.

14. Crosskey, T.D., Beckham, K.S.H., and Wilmanns, M. (2020). The ATPases of the mycobacterial type VII secretion system: Structural and mechanistic insights into secretion. Prog Biophys Mol Biol 152, 25–34. 10.1016/j.pbiomolbio.2019.11.008.

15. Gracy, J., and Chiche, L. (2005). PAT: a protein analysis toolkit for integrated biocomputing on the web. Nucleic Acids Res 33, W65–71. 10.1093/nar/gki455.

16. Nielsen, H. (2017). Predicting Secretory Proteins with SignalP. Methods Mol Biol 1611, 59–73. 10.1007/978-1-4939-7015-5_6.

17. Bagos, P.G., Nikolaou, E.P., Liakopoulos, T.D., and Tsirigos, K.D. (2010). Combined prediction of Tat and Sec signal peptides with hidden Markov models. Bioinformatics 26, 2811–2817. 10.1093/bioinformatics/btq530.

18. Krogh, A., Larsson, B., von Heijne, G., and Sonnhammer, E.L. (2001). Predicting transmembrane protein topology with a hidden Markov model: application to complete genomes. J Mol Biol 305, 567–580. 10.1006/jmbi.2000.4315.

19. Jumper, J., Evans, R., Pritzel, A., Green, T., Figurnov, M., Ronneberger, O., Tunyasuvunakool, K., Bates, R., Žídek, A., Potapenko, A., et al. (2021). Highly accurate protein structure prediction with AlphaFold. Nature 596, 583–589. 10.1038/s41586-021-03819-2.

20. Andreeva, A., Howorth, D., Chothia, C., Kulesha, E., and Murzin, A.G. (2014). SCOP2 prototype: a new approach to protein structure mining. Nucleic Acids Res 42, D310–314. 10.1093/nar/gkt1242.

21. Zhang, Y., and Skolnick, J. (2005). TM-align: a protein structure alignment algorithm based on the TM-score. Nucleic Acids Res 33, 2302–2309. 10.1093/nar/gki524.

22. UniProt Consortium (2021). UniProt: the universal protein knowledgebase in 2021. Nucleic Acids Res 49, D480–D489. 10.1093/nar/gkaa1100.

23. Lew, J.M., Kapopoulou, A., Jones, L.M., and Cole, S.T. (2011). TubercuList--10 years after. Tuberculosis (Edinb) 91, 1–7. 10.1016/j.tube.2010.09.008.

24. Pfam: The protein families database in 2021 - PubMed https://pubmed.ncbi.nlm.nih.gov/33125078/.

25. Szklarczyk, D., Gable, A.L., Nastou, K.C., Lyon, D., Kirsch, R., Pyysalo, S., Doncheva, N.T., Legeay, M., Fang, T., Bork, P., et al. (2021). The STRING database in 2021: customizable protein-protein networks, and functional characterization of user-uploaded gene/measurement sets. Nucleic Acids Res 49, D605–D612. 10.1093/nar/gkaa1074.

26. Anoosha, P., Sakthivel, R., and Gromiha, M.M. (2015). Prediction of protein disorder on amino acid substitutions. Anal Biochem 491, 18–22. 10.1016/j.ab.2015.08.028.

27. Lange, S., Rosenkrands, I., Stein, R., Andersen, P., Kaufmann, S.H.E., and Jungblut, P.R. (2014). Analysis of protein species differentiation among mycobacterial low-Mr-secreted proteins by narrow pH range Immobiline gel 2-DE-MALDI-MS. J Proteomics 97, 235–244. 10.1016/j.jprot.2013.06.036.

28. Talaat, A.M., Lyons, R., Howard, S.T., and Johnston, S.A. (2004). The temporal expression profile of Mycobacterium tuberculosis infection in mice. Proc. Natl. Acad. Sci. U.S.A. 101, 4602–4607. 10.1073/pnas.0306023101.

29. Hauser, M., Mayer, C.E., and Söding, J. (2013). kClust: fast and sensitive clustering of large protein sequence databases. BMC Bioinformatics 14, 248. 10.1186/1471-2105-14-248.

30. MAFFT: a novel method for rapid multiple sequence alignment based on fast Fourier transform - PubMed https://pubmed.ncbi.nlm.nih.gov/12136088/.

